# Essential tremor with tau pathology features seeds indistinguishable in conformation from Alzheimer’s disease and primary age-related tauopathy

**DOI:** 10.1101/2024.10.12.617973

**Authors:** Nil Saez-Calveras, Jaime Vaquer-Alicea, Charles L. White, Yogesh Tak, Stephanie Cosentino, Phyllis L. Faust, Elan D. Louis, Marc I. Diamond

## Abstract

Neurodegenerative tauopathies are characterized by the deposition of distinct fibrillar tau assemblies whose rigid core structures correlate with defined neuropathological phenotypes. Essential tremor (ET) is a progressive neurological disease disease that, in some cases, is associated with cognitive impairment and tau accumulation.. Consequently, we explored the tau assembly conformation in ET patients with tau pathology using cytometry-based tau biosensor assays. These assays quantify tau prion seeding activity present in brain homogenates based on conversion of intracellular tau-fluorescent protein fusions from a soluble to an aggregated state. Prions exhibit seeding barriers, whereby a specific assembly structure cannot serve as a template for a native monomer if the amino acids are not compatible. We recently exploited the tau prion species barrier to define tauopathies by systematically substituting alanine (Ala) in the tau monomer and measuring its incorporation into seeded aggregates within biosensor cells. The Ala scan precisely classified the conformation of tau seeds from diverse tauopathies. We next studied 18 ET patient brains with tau pathology. Only one case had concurrent high amyloid-β plaque pathology consistent with Alzheimer’s disease (AD). We detected robust tau seeding activity in 9 (50%) of the patients. This predominantly localized to the temporal pole and temporal cortex. We examined 8 ET cases with the Ala scan and determined that the amino acid requirements for tau monomer incorporation into aggregates seeded from these ET brain homogenates were identical to those of AD and primary age-related tauopathy (PART), and completely distinct from other tauopathies such as corticobasal degeneration, chronic traumatic encephalopathy, and progressive supranuclear palsy. Based on these studies, tau assembly cores in a pathologically confined subset of ET cases with high tau pathology are identical to AD and PART. This could facilitate more precise diagnosis and therapy for ET patients with cognitive impairment.

## Introduction

Essential tremor (ET) is the most common movement disorder. An estimated 7 million people are affected in the United States representing 2.2% of the entire population ^1,2^. The pathophysiology of ET has not been fully elucidated although there is mounting postmortem evidence that it is neurodegenerative, with postmortem changes observed primarily in the cerebellar cortex ^3,4^. The central clinical feature of ET is an 8-12 Hz kinetic tremor, although a variety of other forms of tremor and motor features are often present ^5,6^. Multiple studies indicate that individuals with ET have poorer cognitive performance than age-matched controls ^7^. Emerging evidence indicates that cognitive problems in ET progress more quickly than in controls ^8,9,10^. Indeed, two prospective, population-based, epidemiological studies, in Madrid and New York, found an association between ET and dementia ^11,12^. In these studies, 11.4% - 25.0% of ET cases (mean age 79.1 - 80.9 years) had prevalent dementia vs. only 6.0% - 9.2% of controls ^11,12^. Furthermore, in both studies, the risk of incident dementia was higher among individuals with baseline ET vs. those without (relative risk [RR] = 1.64 – 1.89) ^11,12^, and tau accumulation in ET was higher than in matched controls^13,14^. The type of tau pathology in ET has not yet been clearly defined from a conformational standpoint, as it includes patterns consistent with Alzheimer’s disease (AD), progressive supranuclear palsy (PSP), and other disorders^13,15,16^.

Brain deposition of tau assemblies defines tauopathies ^17^, and morphological and cellular features have been used to classify these disorders. Recently cryogenic-electron microscopy (cryo-EM) has resolved the atomic structure of *ex vivo* tauopathy filaments, allowing classification of tau assemblies based on their conformation ^18,19^. Cryo-EM reveals distinct tau core filament structures in AD, corticobasal degeneration (CBD), PSP and chronic traumatic encephalopathy (CTE). Importantly, AD and primary age-related tauopathy (PART) uniquely share the same filament structures ^19^, and exhibit similar brain deposition patterns.

Cryo-EM offers atomic-level resolution of structure, yet it is expensive, labor intensive, requires significant tissue mass for extraction, and is biased because only particles chemically extracted and suitable for analysis can be studied. We recently established a rapid and unbiased *in vitro* approach to classify tauopathies from brain homogenates by exploiting tau prion seeding activity and functional genetics. Prions form “strains,” which are distinct amyloid structures that stably propagate *in vivo*, and drive unique patterns of neuropathology and clinical phenotypes. Variation in the amino acid sequence within the native protein recruited to a given assembly creates a “species barrier” with certain strains, such that they will not amplify in the absence of a compatible monomer. This stringency for amino acid sequence protects humans from being infected with prions derived from other species, e.g. cervids or ovines ^20^. Indeed, polymorphism within the prion protein gene (PRNP) has protected individuals from contracting Kuru in New Guinea, where this transmissible prion disorder was endemic ^21,22^.

Based on the concept of the species barrier, we have recently described how systematic alanine (Ala) substitution in the tau monomer (Ala scan), with subsequent measurement of its incorporation into seeded aggregates within biosensor cells, reliably classifies tau seeds derived from diverse tauopathies and correlates specifically with folding of protofilaments (the single protein units of a fibril) ^23^. Without defining structure *per se*, this approach matches the accuracy of cryo-EM in classification of tau protofilaments, without the requirements for extraction and imaging of detergent-insoluble fibrils.

Because of the increased prevalence of cognitive impairment and tau pathology in ET, we determined the underlying tau seed conformation in a subset of ET patients with tau pathology. We used the Ala scan to test whether tau aggregates exhibit a unique conformation. Using cells seeded from ET cases, coupled with Ala scanning for monomer incorporation, we have found that in multiple ET cases the amino acids required for monomer incorporation into aggregates seeded from ET brain are identical to those for AD and PART. This strongly indicates that tau seeds observed in a pathologically confined subset of patients with ET and tau pathology are fundamentally identical to those of AD and PART.

## Materials and Methods

### Human brain samples

#### ET cases

The analyses included brains from 18 ET cases derived from the Essential Tremor Centralized Brain Repository (ETCBR), a longstanding collaboration between investigators at the University of Texas Southwestern Medical Center and Columbia University ^24,25,26^. Established in 2003, the ETCBR banks brains from ET cases throughout the United States. ET diagnoses were carefully assigned by a neurologist specializing in tremor (E.D.L.), using three sequential methods, as has been our practice for 20 years, and as employed in over 50 publications ^3,26, 27^. First, the clinical diagnosis of ET was initially assigned by the treating neurologists, and second, confirmed by E.D.L. using semi-structured clinical questionnaires, medical records, and Archimedes spirals with the following criteria: (i) moderate or greater amplitude kinetic tremor (rating of 2 or higher ^28^) in at least one of the submitted Archimedes spirals; (ii) no history of PD or dystonia; and (iii) no other etiology for tremor (e.g., medications, hyperthyroidism) ^27,26^. Third, a detailed, videotaped, neurological examination was performed, from which action tremor was rated and a total tremor score assigned (range 0-36 [maximum]) ^27,26^. These data were used to assign a final diagnosis of ET ^27,26^, reflecting published diagnostic criteria [moderate or greater amplitude kinetic tremor (tremor rating ≥ 2) during three or more activities or a head tremor in the absence of PD or other known causes] ^28^, of demonstrated reliability and validity ^29,30^. No ET cases reported a history of traumatic brain injury, exposure to medications associated with cerebellar toxicity (e.g., phenytoin, chemotherapeutic agents), or heavy ethanol use ^31^. Every 6-9 months, a follow-up semi-structured telephone evaluation was performed, and hand-drawn spirals were collected; a detailed, videotaped, neurological examination was repeated if there was concern about a new, emerging movement disorder.

#### Other tauopathy cases

AD, PART, CBD, and PSP human brain tissue was obtained through the University of Texas Southwestern Medical Center Neuropathology Brain Bank with Institutional Review Board (IRB) approval. The CTE case was obtained from CTE Center at Boston University (Adapted from Vaquer-Alicea, *bioRxiv 2024* ^23^).

### Tissue processing and neuropathological examination

Brains from the New York Brain Bank had a complete neuropathological assessment with standardized measurements of brain weight and postmortem interval (hours between death and placement of brain in a cold room or upon ice) ^29,30^. Standardized blocks were harvested from each brain and processed, and 7 μm-thick formalin-fixed paraffin-embedded sections were stained with Luxol fast blue/hematoxylin and eosin (LH&E) ^32,33^. Additionally, selected sections were stained by the Bielschowsky method, and with mouse monoclonal antibodies to phosphorylated tau (clone AT8, Research Diagnostics, Flanders, NJ) and β-amyloid (clone 6F/3D, Dako, Carpenteria, CA) ^32,33^. All tissues were examined microscopically by a senior neuropathologist blinded to clinical information ^32,33^. Alzheimer’s disease staging for neurofibrillary tangles (Braak NFT) ^34^, Consortium to Establish a Registry for Alzheimer’s disease (CERAD) ratings for neuritic plaques ^35,36^ and Thal β-amyloid stages were assigned ^37,38^. For Braak NFT staging, stages I and II indicated NFTs confined to the entorhinal region, III and IV limbic region involvement (i.e. hippocampus), and V and VI moderate to severe neocortical involvement ^34^. An additional classifier (PSP) was added for subjects with PSP-like pathology ^39^. The level of AD neuropathologic change (ADNC) was rated according to National Institute on Aging and Alzheimer’s Association (NIA-AA) guidelines using an “ABC” score as none, low, intermediate or high ^38^.

One ET case with pathologically confirmed high ADNC was examined in this study as a positive control for detection of AD-type tau in seeding assays (Table 1, case 1). The remaining ET cases were chosen based on the following criteria: 1) presence of no (Thal A0, CERAD C0) or few amyloid deposits (Thal A1 or CERAD C1); 2) higher Braak NFT stage (5 – 6), reflecting at least some extension into frontal association cortices and/or temporal pole; and/or 3) presence of PSP-type changes. Frozen samples from cerebral cortex regions including frontal (BA9), motor (BA4), parietal (BA7), occipital (BA31), temporal (BA37), and temporal pole (BA38) were assayed according to sample availability. In cases with PSP-type changes detected, cerebellar cortex and dentate nucleus were assayed, as available. Cases that had only focal PSP-type pathology not meeting current Rainwater criteria ^39^ were designated as “PSP-like.”

### Tissue homogenization

Frozen brain was suspended in tris-buffered saline (TBS) containing cOmplete mini protease inhibitor tablet (Roche) at a concentration of 10% weight/volume. Homogenates were prepared by probe homogenization using a Power Gen 125 tissue homogenizer (Fisher Scientific) in a vented hood. The homogenates were then centrifuged at 21,000 x g for 20 min. The supernatant was collected as the total soluble protein lysate. Protein concentration was measured using a Pierce 660 assay (Pierce). Fractions were aliquoted into low-binding tubes (Thermo Fisher) and frozen at -80°C until future use.

### Tau RD biosensor cell lines

The following tau repeat domain (RD) biosensor cell lines were used for tau seeding assays in this study:

#### Tau RD 3R-Cer/4R-Ruby (3R/4R) cells

HEK293T cells containing the 3-repeat (3R) version of the wild-type human tau RD (residues 246-408) C-terminally fused to mCerulean (Cer), and the 4R version fused to mRuby fluorescent protein (Rub) were used for tau seeding assays and the 3R/3R tau Ala scan. These biosensor cells exhibit high seeding after exposure to brain homogenates from certain tauopathies (e.g. AD) but not others (e.g. CBD) ^23^.

#### Tau RD 4R-Cer/4R-Ruby (4R/4R) cells

HEK293T cells containing the 4R version of the wild-type human tau RD (residues 246-408), C-terminally fused to monomeric Cerulean3 (Cer) and monomeric Ruby fluorescent protein (mRuby3) were used for tau seeding assays and the 4R/4R tau Ala scan. These biosensor cells exhibit high seeding after exposure to brain homogenates from certain tauopathies (e.g. CBD, PSP) but not others (e.g. AD) ^23^.

### Tau (246-408) Ala scan library

The WT-tau (246-408)-Ala mutant library used in this study was previously generated and described in detail ^23^. Twist Biosciences synthesized gene fragments encoding the human tau sequence from residues 246 to 408 with Ala codon substitutions (GCC) at each position. These gene fragments were then conjugated to a common lentiviral plasmid harboring the mEOS3.2 fluorescent protein, thus generating an arrayed library of plasmids with the different tau Ala mutants fused to mEOS3.2. All plasmids were verified by Sanger sequencing.

### Biosensor cell transduction with brain homogenates and tau seeding assay

*Tau RD 3R-Cer/4R-Ruby (3R/4R)* and *Tau RD 4R-Cer/4R-Ruby (4R/4R)* cells were plated at a density of 20,000 cells per well of a 96-well plate. 24 hours later the cells were transduced with a complex of 10 μg of 10% w/v brain homogenate, 0.75 μl of lipofectamine 2000 (Invitrogen) and 9.25μl of Opti-MEM for a final treatment volume of 20 μl per well. 72 hours after transduction, cells were harvested using 0.25% trypsin digestion for 5 min at 37 °C, quenched with Dulbecco’s Modified Eagle Medium (DMEM), and transferred to 96-well U bottom plates. They were then centrifuged for 5 minutes at 1500 rpm, the supernatant was aspirated, and the cells were fixed in 2% paraformaldehyde in PBS for 10 minutes. The cells were then centrifuged again and resuspended in 150 μl of PBS for subsequent analysis using flow cytometry.

### Ala scan

For the Ala scan incorporation assay, 3R/4R or 4R/4R cells were plated at 20,000 cells per well in 96-well plates and treated with sonicated brain homogenates as described above. After 48 hours, the cells were replated at a concentration ratio of 1:6 into 6 new 96 well plates to generate technical replicates. The cells were then treated with the library of WT-tau (246-408)-Ala mutant lentivirus to achieve sufficient transduction efficiency (20 μl of media containing lentivirus). One WT tau Ala mutant was added per well. 72 hours after transduction the cells were harvested and fixed in paraformaldehyde as described above (See *Biosensor cell transduction with brain homogenates and tau seeding assay*).

### Flow cytometry

The BD LSRFortessa flow cytometer was used to perform Fluorescence Resonance Energy Transfer (FRET) analysis. To measure CFP or Cer signal, and FRET, cells were excited with the 405 nm laser, and fluorescence was captured with a 488/50 nm and 525/50 nm filter. To measure YFP and mEOS3.2, cells were excited with a 488 nm laser and fluorescence was captured with a 525/50 nm filter. To measure Ruby, cells were excited with a 561 nm laser and fluorescence was detected using a 610/20 nm filter.

### Data analysis

Seeded 3R/4R or 4R/4R biosensor cells transduced with the WT tau Ala scan library were analyzed by gating for homogeneous side-scatter and forward-scatter. The FRET signal between Cer and Ruby, measured in the Pacific Blue and Qdot605 channels, was used to gate for cells that contained seeded aggregates in these biosensors. Within this population, a narrow gate for cells positive for FITC (non-photoconverted mEos3.2) was selected, within which FRET between Cer and mEOS3.2 was gated. Within this population, a population of bright cells were selected to calculate the median fluorescence intensity (MFI) for the AmCyan channel. All gates were kept constant across the incorporation assay and across technical replicates. The fluorescence intensity of the FRET signal in the AmCyan channel was subsequently used for the incorporation assay analysis.

### Data processing

FRET median fluorescence intensity (MFI) values between Cerulean and mEOS3.2 measured in the AmCyan channel were normalized by plate to prevent batch variation. Within our analysis, most Ala residues in the N- and the C-termini of the sequence did not impact incorporation. Thus, we used these values, and the lowest value in each plate to normalize the data, following this formula:

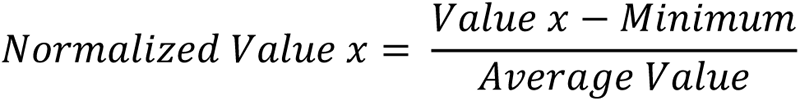

where *Value x* is the FRET MFI for each position*, Minimum* is the minimum FRET value in the scan, and *Average Value* is the Average MFI for the first 20 residues (on the first plate) or the last 10 residues (on the second plate). The average of three technical replicates was used for downstream analysis.

## Results

### Case characteristics

We studied 18 ET cases (Table 1). One exhibited prominent Aβ pathology and high Braak NFT stage, leading to a postmortem diagnosis of high ADNC (case 1). The other 17 cases consisted of cases with high NFT burden (Braak NFT ≥3) or PSP findings, and low-to-absent Aβ pathology as assessed by Thal and CERAD staging. These cases had received additional postmortem diagnoses of typical PSP, combined PSP and PART.

**Table 1:** Demographic and neuropathologic data.

| **Table 1: Demographic and neuropathologic data** |  |  |  |  |  |  |
| --- | --- | --- | --- | --- | --- | --- |
| **Case** | **Age (Range)** | **Sex** | **Neuropathologic diagnosis(es)** | **Thal** | **Braak NFT** | **CERAD** |
| 1 | 86-90 | F | ADNC, high | A3 | 6 | 3 |
| 2 | 96-100 | M | PSP, combined, mild; Infarcts, cortico-subcortical, old, left pre-frontal and right frontal | A1 | 6 | 0 |
| 3 | 91-95 | F | PSP, combined, moderate; Infarct, old, large, territory of the anterior cerebral artery, left | A0 | 6 | 0 |
| 4 | 71-75 | F | PSP, combined, mild; ADNC, low | A1 | 5 | 1 |
| 5 | 91-95 | F | PSP, combined, mild; ADNC, low | A1 | 3 | 1 |
| 6 | 101-105 | M | PSP, typical, mild | A1 | 5 | 0 |
| 7 | 91-95 | M | PART; PSP-type changes, rare | A0 | 6 | 0 |
| 8 | 86-90 | F | PART | A0 | 5 | 0 |
| 9 | 91-95 | F | PSP, typical, mild | A0 | 3 | 0 |
| 10 | 81-85 | F | PART; PSP-type changes, rare | A1 | 4 | 0 |
| 11 | 96-100 | M | PSP, combined, moderate*; ADNC, low | A1 | 6 | 0 |
| 12 | 86-90 | F | PSP, typical, mild | A1 | 5 | 0 |
| 13 | 86-90 | F | ADNC, low [possible PART] | A1 | 5 | 1 |
| 14 | 86-90 | M | PART | A0 | 5 | 0 |
| 15 | 71-75 | M | PART | A0 | 5 | 0 |
| 16 | 86-90 | F | ADNC, low [possible PART] | A1 | 5 | 1 |
| 17 | 76-80 | F | PART | A0 | 5 | 0 |
| 18 | 81-85 | F | ADNC, low [possible PART] | A1 | 5 | 0 |
| ADNC, Alzheimer Disease Neuropathologic Changes; Braak NFT (Braak staging for Neurofibrillary Tangles); CERAD (Consortium to Establish a Registry for Alzheimer's disease); PART, primary age-related tauopathy; PSP, progressive supranuclear palsy. *, PSP changes were more prominent in subcortical regions in this case. |  |  |  |  |  |  |

**Table 2:** Regions used for Tau seeding analysis.

| **Table 2: Regions used for Tau seeding analysis** |  |  |  |  |  |  |  |  |  |  |
| --- | --- | --- | --- | --- | --- | --- | --- | --- | --- | --- |
| **Case** | **Age (Range)** | **Sex** | **TC (BA37)** | **TP (BA38)** | **FC (BA9)** | **MC (BA4)** | **PC (BA7)** | **OC (BA31)** | **Cbl Ctx** | **DN** |
| 1 | 86-90 | F | X |  |  |  |  |  |  |  |
| 2 | 96-100 | M | X |  |  |  |  |  | X |  |
| 3 | 91-95 | F | X |  |  |  |  |  | X | X |
| 4 | 71-75 | F | X |  |  |  |  |  | X | X |
| 5 | 91-95 | F | X |  |  |  |  |  | X |  |
| 6 | 101-105 | M | X |  |  |  |  |  | X | X |
| 7 | 91-95 | M | X | X | X | X | X | X | X |  |
| 8 | 86-90 | F | X | X | X |  | X | X |  |  |
| 9 | 91-95 | F | X | X | X | X | X | X | X | X |
| 10 | 81-85 | F |  | X | X | X | X | X | X | X |
| 11 | 96-100 | M | X | X | X | X | X | X | X | X |
| 12 | 86-90 | F |  | X | X | X | X | X | X | X |
| 13 | 86-90 | F |  | X | X |  | X | X |  |  |
| 14 | 86-90 | M |  | X | X |  |  | X |  |  |
| 15 | 71-75 | M | X | X | X |  | X | X |  |  |
| 16 | 86-90 | F |  | X | X |  | X | X |  |  |
| 17 | 76-80 | F |  | X |  |  | X | X |  |  |
| 18 | 81-85 | F | X |  |  |  |  |  |  |  |
| TC: Temporal Cortex; TP: Temporal Pole; FC: Frontal Cortex; MC: Motor Cortex; PC: Parietal Cortex; OC: Occipital Cortex; Cbl Ctx: Cerebellar Cortex; DN: Dentate Nucleus |  |  |  |  |  |  |  |  |  |  |

### ET cases with high NFT burden exhibit tau seeding activity

We first tested whether ET brain homogenates (Table 2) from multiple brain regions seeded 3R/4R (as observed in AD or PART), or 4R/4R biosensors (as observed in CBD, CTE, and PSP) (Fig. 1S). Multiple cases (9/18, 50%) exhibited robust seeding activity at or above 5% FRET positive cells predominantly on 3R/4R biosensors. In these cases, the temporal cortex and temporal pole regions scored highest (cases 1, 7, 11-15, 17) (Fig. 1A). In case 16, the parietal cortex scored highest. For some we observed equivalent seeding on 3R/4R and 4R/4R biosensors (cases 1, 7, 12-14) (Fig. 1B). Given that ET patients exhibit degenerative changes in and around Purkinje cells in the cerebellar cortex ^25,40,41,42^ we tested cerebellar cortex and dentate nucleus, which had low or absent seeding. The other ET cases (2,3,4,5,6,8,9,10,18) exhibited no/low tau seeding (below 5%) (Fig. 1S).

**Figure 1:**
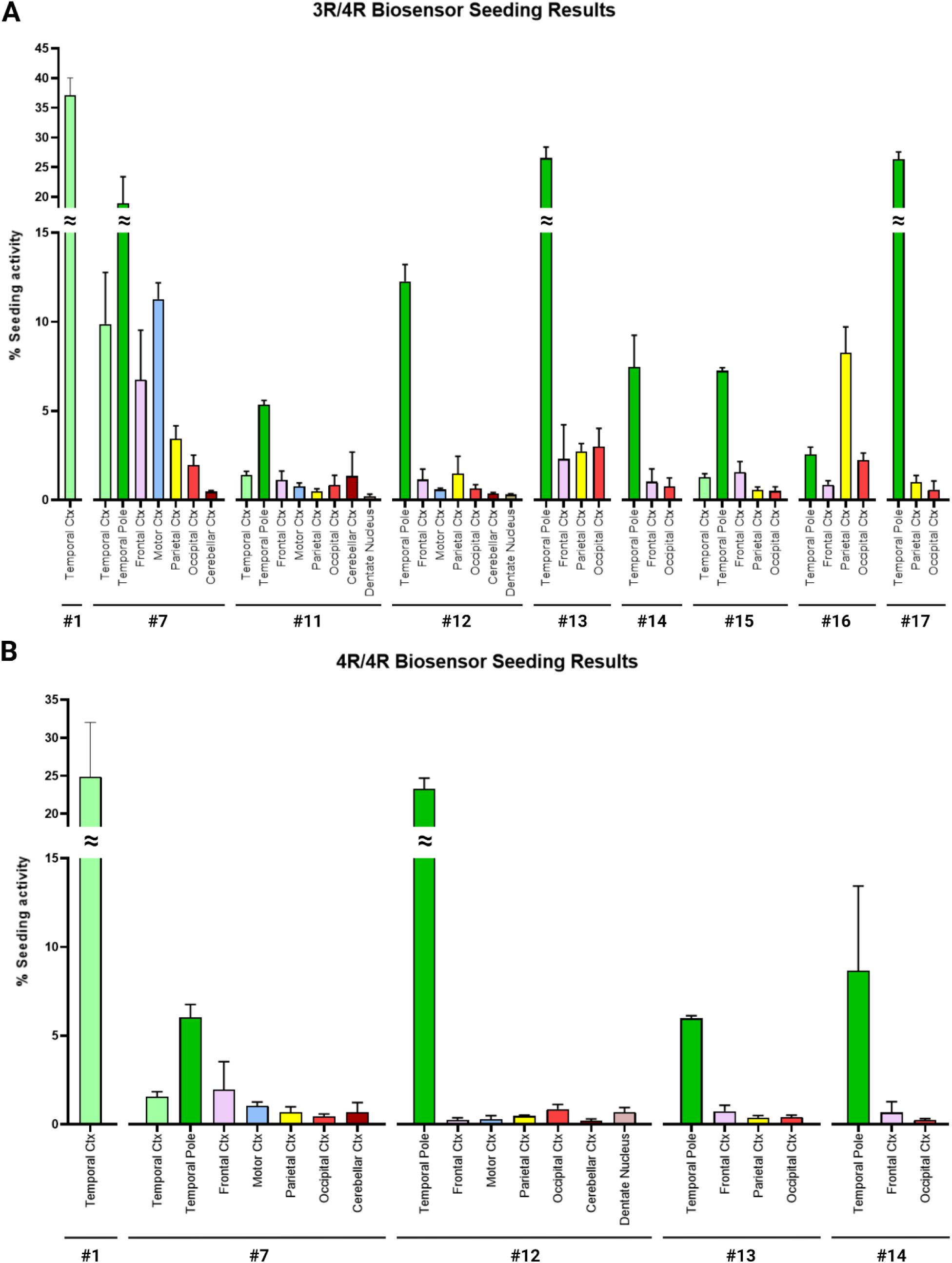
ET cases exhibit seeding in tau biosensor cells. Brain homogenates from ET patients were transduced into 3R/4R (**A**) and 4R/4R (**B**) biosensor cells. ET cases with tau seeding are reported. Strong seeding was recorded primarily in the temporal pole and the temporal cortex, except in case 16 (parietal cortex). Cases 1, 7, 12-14 had similar seeding in 3R/4R and 4R/4R biosensors. *Ctx:* Cortex.

### Ala scan in ET cases is identical to AD and PART

To evaluate the conformation of the tau assemblies in ET cases, which varied widely in their postmortem diagnoses, we used the recently validated Ala scan. We analyzed 8 ET cases where sufficient tissue with seeding activity was available (Table 3). Case #16 could not be completed due to insufficient tissue availability despite having >5% seeding on 3R/4R biosensors. The Ala scan is based on a two-step process in which brain homogenates are seeded into appropriate biosensors containing 3R/4R or 4R/4R tau RD, followed by exposure to lentiviruses individually expressing 4R tau monomer with Ala substitutions through the RD (Fig. 2). Prior data indicated that tauopathies may be discriminated in part by whether they seed more efficiently onto 3R/4R biosensors (e.g. AD) vs. 4R/4R biosensors (e.g. PSP, CBD)^23,43,44^. The effect of each Ala point substitution on the incorporation of tau monomers into seeded aggregates serves as a conformational ’fingerprint’ of the underlying tau aggregate. This fingerprint can accurately identify the specific type of tauopathy. ^23^.

**Table 3:** Cases Selected for 3R/4R Ala scan.

| **Table 3: Cases Selected for 3R/4R Ala scan** |  |  |  |  |
| --- | --- | --- | --- | --- |
| **Case #** | **Age (Range)** | **Sex** | **Seeding on 3R/4R** | **Region for 3R/4R Ala scan** |
| 1 | 86-90 | F | Yes | temporal cortex |
| 7 | 91-95 | M | Yes | temporal pole |
| 11 | 96-100 | M | Yes | temporal pole |
| 12 | 86-90 | F | Yes | temporal pole |
| 13 | 86-90 | F | Yes | temporal pole |
| 14 | 86-90 | M | Yes | temporal pole |
| 16 | 86-90 | F | Yes | parietal cortex |
| 17 | 76-80 | F | Yes | temporal pole |
| *\*An Ala scan was not completed for Case #15 despite >5% seeding due to insufficient tissue* |  |  |  |  |

**Figure 2:**
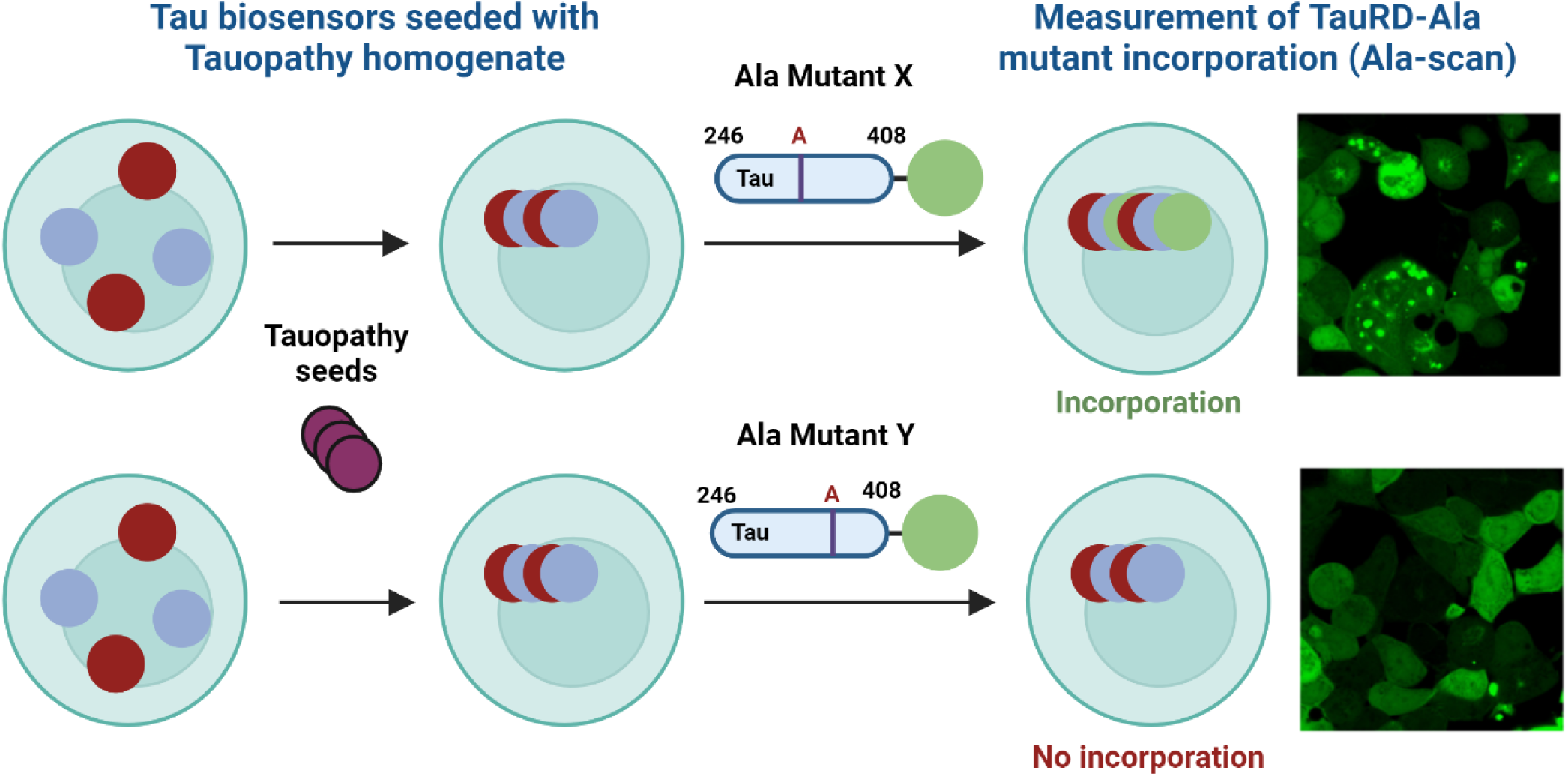
Ala scan diagram. Biosensor cells (i.e. 3R/4R or 4R/4R) were transduced with brain homogenate. Cells were then transduced with lentivirus containing a library of tau RD Ala point mutants (e.g. Mutant X, Mutant Y). The differential incorporation of monomer containing these point mutants into the aggregates is measured by FRET.

To confirm the Ala scan fidelity, we scanned individual cases of PART and AD, which feature identical cryo-EM filament structures ^23^. The PART and AD cases preferentially seeded on 3R/4R biosensors and their profiles were indistinguishable (Fig. 3). This contrasted with the profiles observed for other tauopathies (CTE, PSP, CBD). In each case, we observed Ala scans concordant with our prior published work ^23^.

**Figure 3:**
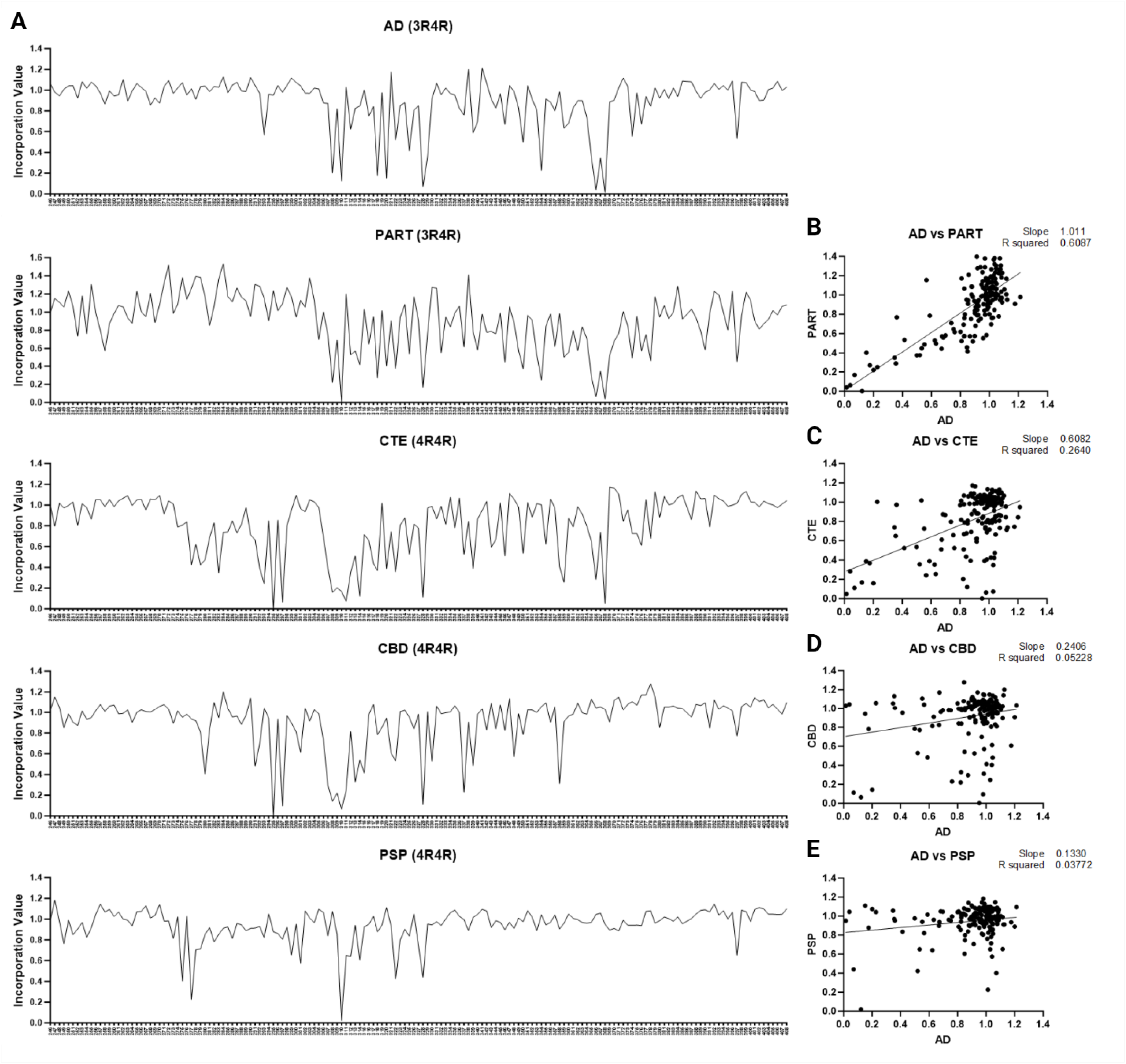
Ala scan for AD, PART, CTE, PSP and CBD cases. AD, PART, CTE, CBD and PSP case brain homogenates were incubated with 3R/4R (AD, PART) or 4R/4R (CTE, PSP, CBD) biosensors based on preferential seeding. After 48 hours (to allow inclusion formation) cells were plated in triplicate and incubated with the WT-Tau (246-408)-Ala mutant lentivirus library. (**A**) Line plots of the tau Ala scans for AD, PART, CTE, CBD and PSP. (**B**) AD and PART scans were highly correlated. AD scans did not correlate with (**C**) CTE; (**D**) CBD; (**E**) PSP.

All ET homogenates seeded on 3R/4R biosensors. To obtain signal sufficient for the 3R/4R Ala scan, we selected brain regions with the highest level of seeding. We used temporal cortex of case 1; the temporal pole of cases 7, 11-14, 17; and the parietal cortex of case 16. The Ala scan of all ET cases was identical to AD and PART (Fig. 4). To assess this in an unbiased fashion we used cluster analysis to group cases according to similarity (Fig. 5). Confirming their similarity, the AD and PART cases were admixed with the ET cases, and distinct from other tauopathies. The CTE case exhibited a partial correlation with these cases, consistent with the structural similarity of the CTE vs. AD and PART protofilament folds ^19,23^. We observed poor correlation with the CBD and PSP cases (Fig. 5). To compare the residue hits across cases, we used heat map analysis, which also highlighted the similarities between ET, AD, and PART, with hits at positions I308, Y310, V318, S320, I328, I354, G366 and N368 (Fig. 6).

**Figure 4.**
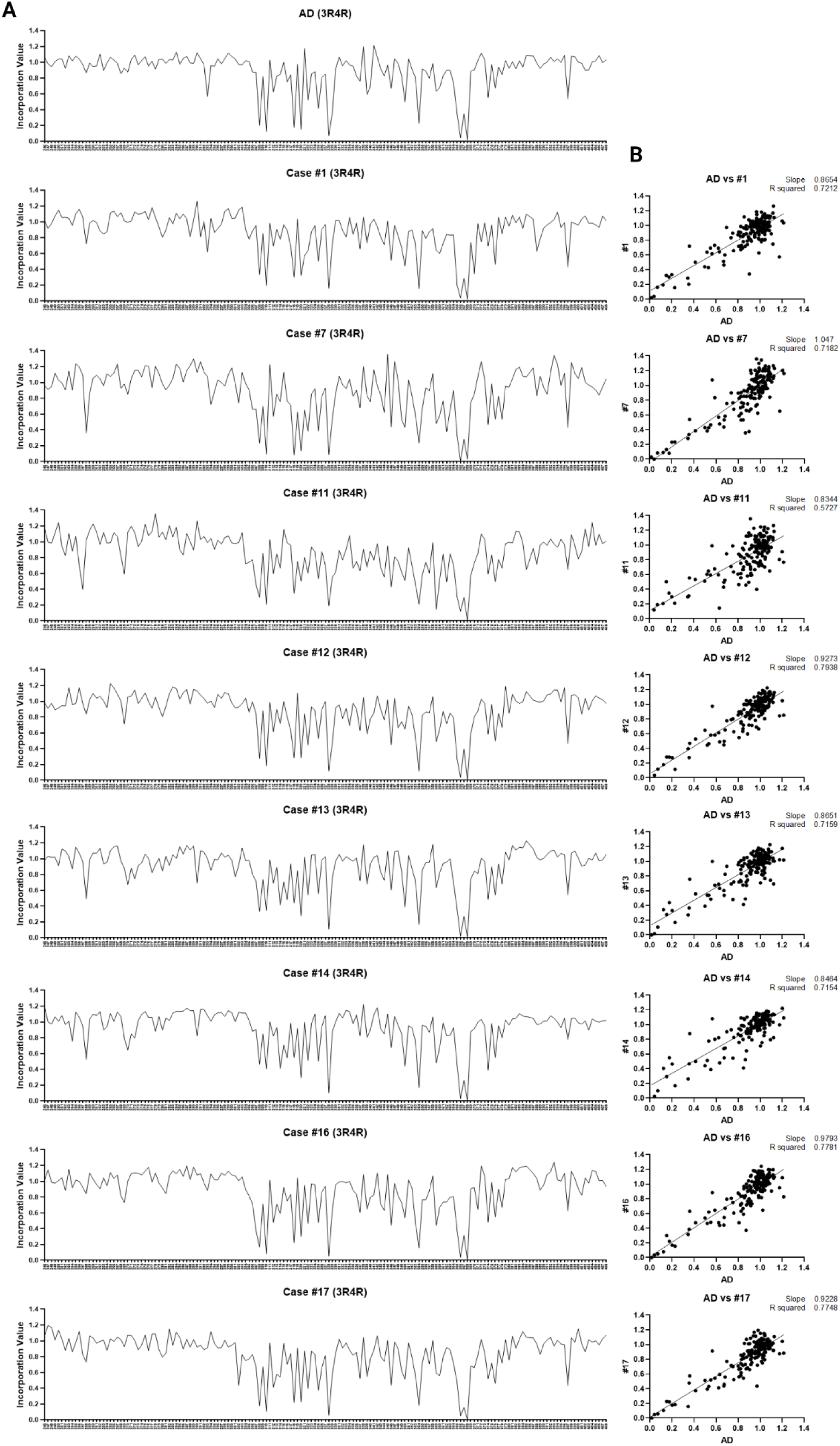
3R/4R Ala scan of ET cases. Brain homogenates from the highest seeding regions (cases 1, 7, 11-14, 16) were transduced into 3R/4R biosensors. After 48 hours, the cells were incubated with the WT-Tau (246-408)-Ala mutant lentivirus library. (**A**) Line plots of the 3R/4R Ala scan replicates indicate effects of substitution for each residue (dot) in the setting of AD vs. ET cases. (**B**) Each ET scan correlated highly with the AD 3R/4R Ala scan.

**Figure 5.**
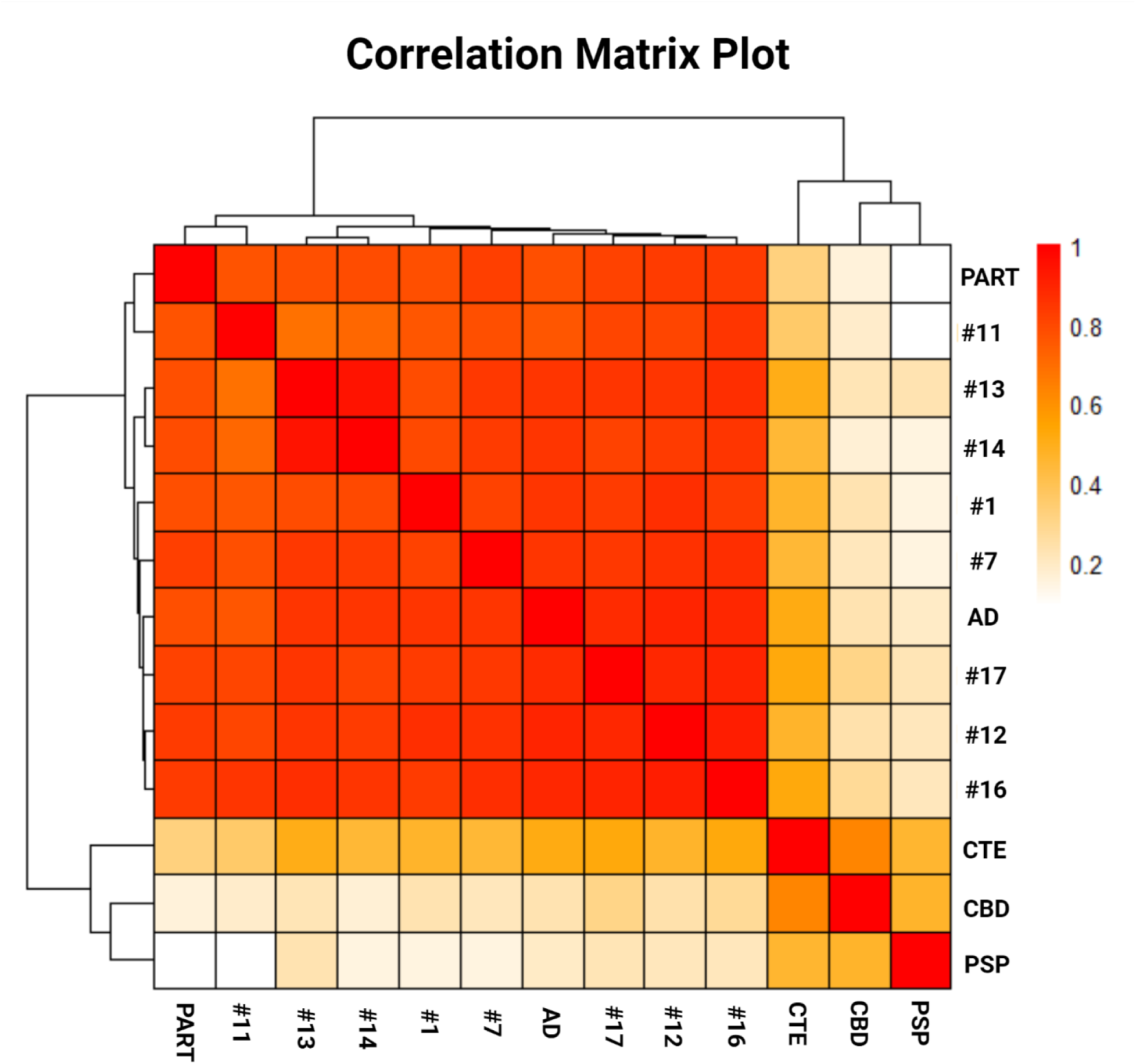
Clustering of ET, AD, PART, CTE, CBD, PSP cases based on the Ala scan. Cluster analysis of Ala scans for all ET cases (3R/4R) reveals grouping of these cases with admixed AD and PART (3R/4R). CTE, CBD, and PSP cases (4R/4R) are poorly correlated with the ET cases.

**Figure 6.**
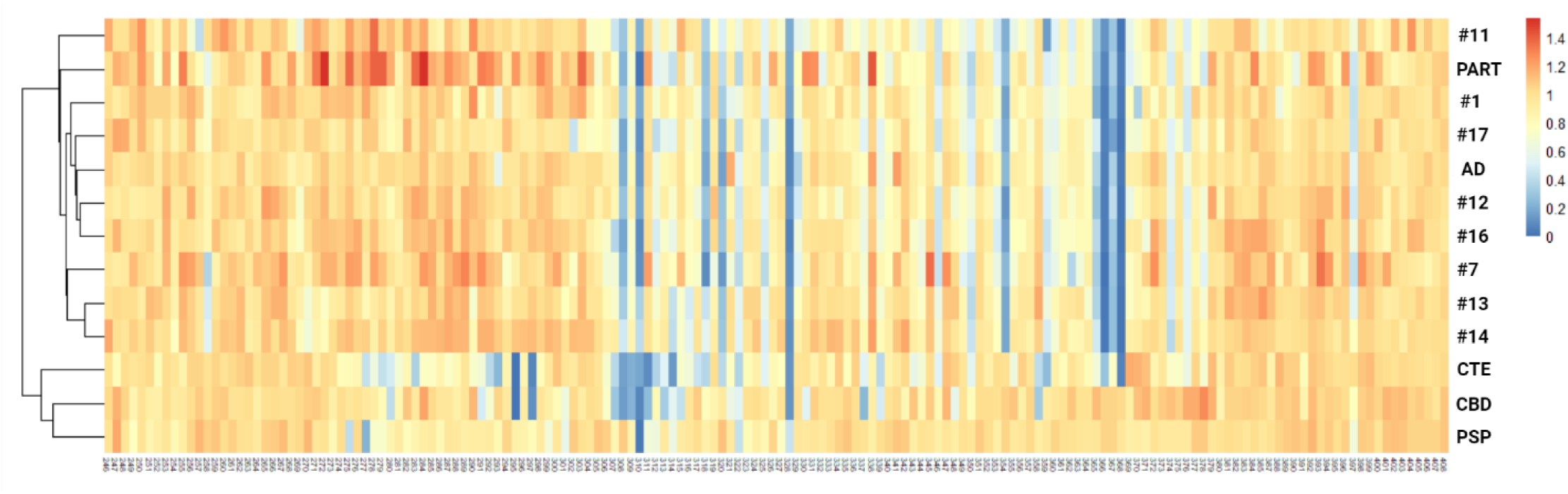
Heatmap incorporation profile of the 3R/4R Ala scan for ET, AD, PART, CTE, CBD, PSP. Heatmap of the 3R/4R Ala scan indicates similar tau Ala hits in ET, AD and PART cases at residue positions I308, Y310, V318, S320, I328, I354, G366, N368.

As mentioned above, five ET cases exhibited seeding (≥5%) on 4R/4R biosensors in addition to the 3R/4R biosensors (cases 1, 7, 12-14). As an additional analysis (Fig. 7), we performed a 4R/4R Ala scan of these cases. For comparison, we also seeded an AD case on 4R/4R biosensors, where we were able to extract a profile, despite lower seeding efficiency compared to 3R/4R (Fig. 7). Given that cases 7 and 12 had a neuropathological diagnosis of PSP, we also compared their profiles to the 4R/4R Ala scan of PSP. The results showed that, despite lower resolution than the 3R/4R Ala scan, the 4R/4R Ala scan profile of these ET cases resembled the pattern observed for AD. Meanwhile, the cases did not match the PSP profile.

**Figure 7:**
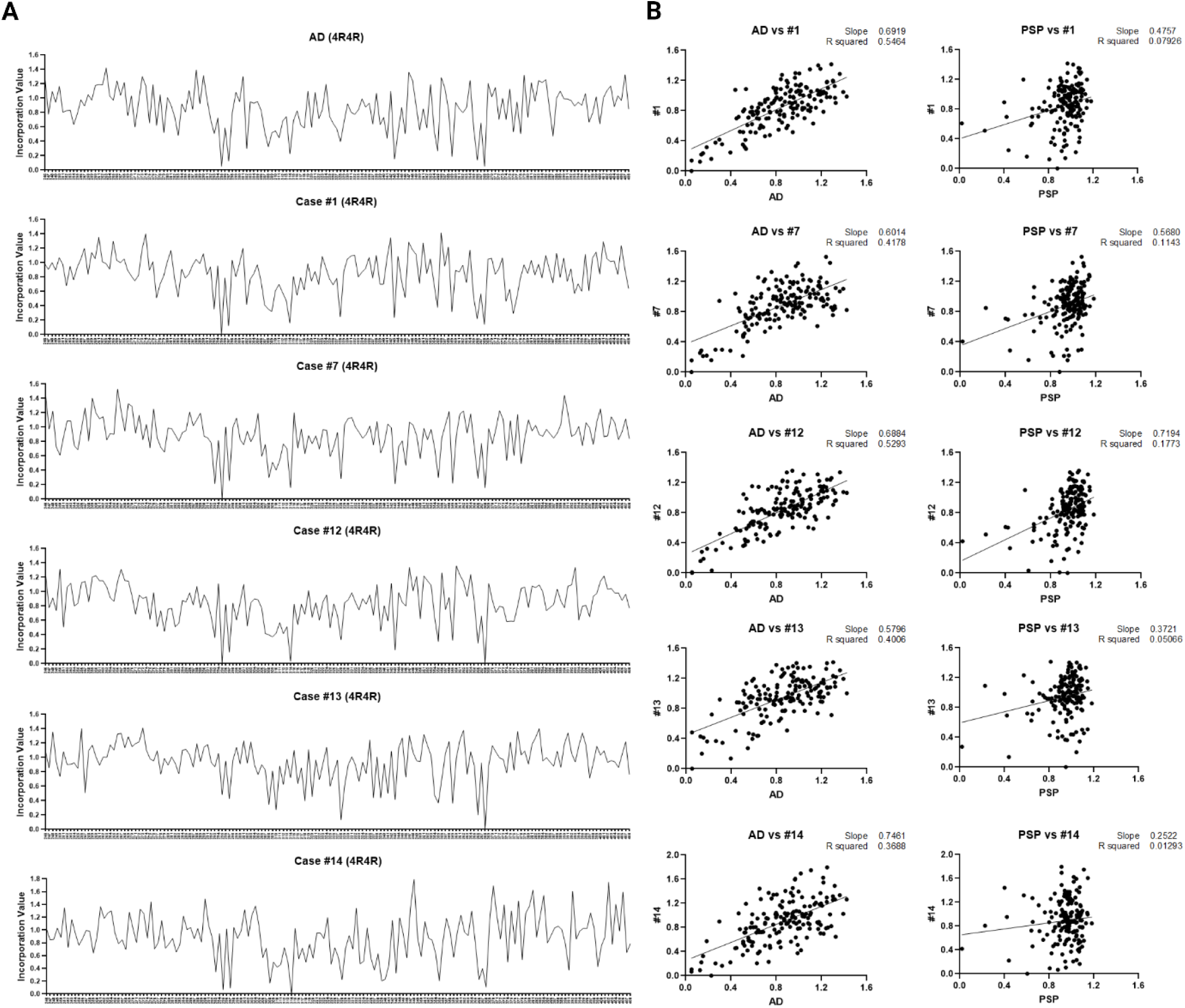
4R/4R Ala scan of ET cases. Brain homogenates from the highest seeding regions for cases 1, 7, 12, 13, 14 and an AD case were transduced into 4R/4R biosensors. After 48 hours, the cells were incubated with the WT-Tau (246-408)-Ala mutant lentivirus library. (**A**) Line plots of the 4R/4R Ala scan indicate the effects of substitution for each residue in AD and in the ET cases. (**B**) Scatter plot comparison between each ET case 4R/4R Ala Scan and the AD and PSP 4R/4R Ala Scans. The plots showed a good correlation with AD but not with PSP.

## Discussion

ET is recognized as a neurodegenerative disease primarily characterized by cerebellar cortical degeneration ^25,40,45,46^. A notable and poorly understood aspect of ET is the increased risk for developing Lewy body ^47^ and tau pathologies ^13,15,16^. Clinically, this translates into a substantially higher risk for PD ^48^ and dementia ^12^ in ET patients compared to those without ET. It is well known that ET cases exhibit a higher pathological tau burden and cognitive impairment when compared to age-matched controls ^13,15,16^. Our study revealed that in ET cases where tau seeding was present, the tau seed conformation assessed by the Ala scan matched the profile in AD and PART, and differed from other tauopathies such as CBD, CTE, and PSP. This suggests that a subset of ET cases with tau pathology harbors tau assemblies indistinguishable from those in AD and PART.

### A subset of ET patients exhibit tau seeding

Experimental and observational evidence suggests that progression and diversity of neurodegenerative tauopathies is based on prion mechanisms, whereby a pathological assembly that forms in one cell can escape, enter a connected cell, and serve as a template for its own replication ^43^. This process has been replicated in simple biosensor cell models such as were used here ^49,50^. This study is the first to identify tau seeding across multiple brain regions in ET patients with tau pathology, indicating that a subset of these patients feature tau assemblies capable of propagation. Tau seed deposition and propagation may explain the origin and progression of cognitive impairment in ET ^51,52,53,54^. Our findings are supported by prior neuropathological studies which revealed that ET patients have a tau NFT burden greater than expected for age ^13,55^, including in cognitively normal cases ^13^.

### Tau assemblies in ET have conformational characteristics of AD and PART

We have recently determined that the amino acid requirements for monomer incorporation into a pre-existing tau aggregate seeded by brain homogenate (Ala scan) precisely classifies tauopathies ^23^. Here we determined that 8 ET brain homogenates exhibited Ala scans identical to AD and PART, and completely distinct from other major tauopathies. Without cryo-EM analysis, the current state-of-the-art, it is not possible to determine the structure of tau assemblies in ET, but the Ala scan is an unbiased, functional genetic analysis with proven power. Importantly, given the low requirements for brain tissue (approximately 1000 µg/scan), it can be applied in situations where a sufficient mass of tau assemblies may not be extractable for cryo-EM. Based on the conformation of tau seeds, our findings place ET patients with tau pathology in the same pathological category as AD and PART^19^. While AD typically does not seed efficiently onto 4R/4R biosensors, we observed a subset of ET cases that efficiently seeded both 3R/4R and 4R/4R biosensors (cases 1, 7, 12-14). However, when we performed a 4R/4R Ala scan analysis of these cases, the profile matched that observed for the 4R/4R Ala scan in AD, although the seeding efficiency for these cases in these biosensors was higher than that of AD. While Ala scans were identical, this could indicate that the tau isoform composition in some ET cases differs from that of AD and PART ^15^, as 4R tau aggregates typically seed efficiently onto 4R/4R biosensors. In ET, seeding primarily localized to the temporal pole and temporal cortex, matching a pattern that can be observed in AD and PART. However, in one case seeding was primarily located in the parietal cortex (case 16), and in three cases it was detected in the occipital cortex (cases 7, 13, 16), with minimal or absent Aβ pathology. This pattern is atypical for the distribution of NFT pathology typically observed in PART or AD ^56^, and may be a neuropathological indicator that the tau isoform composition varies in some of these ET patients.

Notably, although some of the cases analyzed had a neuropathological diagnosis of PSP (cases 7, 11, 12), the Ala scan profiles for these cases did not match that of PSP. Notably in cases 7 and 12 we completed an Ala scan in 3R/4R and in 4R/4R biosensors, and in neither of them did the profile resemble PSP. However, one must consider that the area used for analysis in this study (temporal pole/temporal cortex) was not the optimal to detect PSP-type changes. The presence of AD and PSP co-pathology has been described in advanced dementia ^57^. Given this, we cannot rule out that a PSP fold may be present in other regions that have not been sampled in this study. Thus, further assessment with sampling of additional regions will be necessary.

### Study limitations

One key limitation of our study is that it was restricted to ET patients with tau pathology. Therefore, these findings cannot be generalized to all patients with ET, but only to those in which tau pathology with seeding is present. A second limitation is the restricted access to different brain regions. For some of the patients the temporal pole/cortex were the only cortical regions available for analysis. As mentioned, this may have limited the sensitivity to detect a PSP tau conformation in these patients. Future studies with more comprehensive regional sampling will help us better understand the spatial distribution of tau seeding in ET and compare it to AD and PART. Further analysis of insoluble tau isoform composition in ET will help clarify the proportion of tau isoforms involved in the pathological assemblies.

### Clinical implications for ET patients with tau pathology

Our findings suggest new diagnostic and therapeutic avenues for ET with tau pathology and/or cognitive impairment, because a subset of cases appear to have the same underlying tau assembly conformation as AD and PART. For example we predict that tau positron emission tomography (PET) imaging radiotracers that are effective in AD ^58,59,60^ will also identify ET patients with tau pathology ^61^. Similarly, if anti-tau therapies work in AD, they might ultimately also be effective in ET patients with tau pathology ^62^. Taken together, our findings have profound implications for more impactful care of ET patients.

## Author contributions

Conceptualization: NSC, EDL, MID

Methodology: NSC, JVA, PLF, CLW

Investigation: NSC, JVA, YT, PLF, SC

Visualization: NSC, JVA, PLF, EDL, MID

Supervision: EDL, MID

Writing - original draft: NSC, EDL, MID

Writing - reviewing and editing: NSC, MID, JVA, PLF, EDL

## Data and materials availability

All data are available without any restriction upon request to the authors or is already available in the main text and supplement.

