## Supplementary Figure 1S for "Essential tremor with tau pathology features seeds indistinguishable in conformation from Alzheimer’s disease and primary age-related tauopathy"

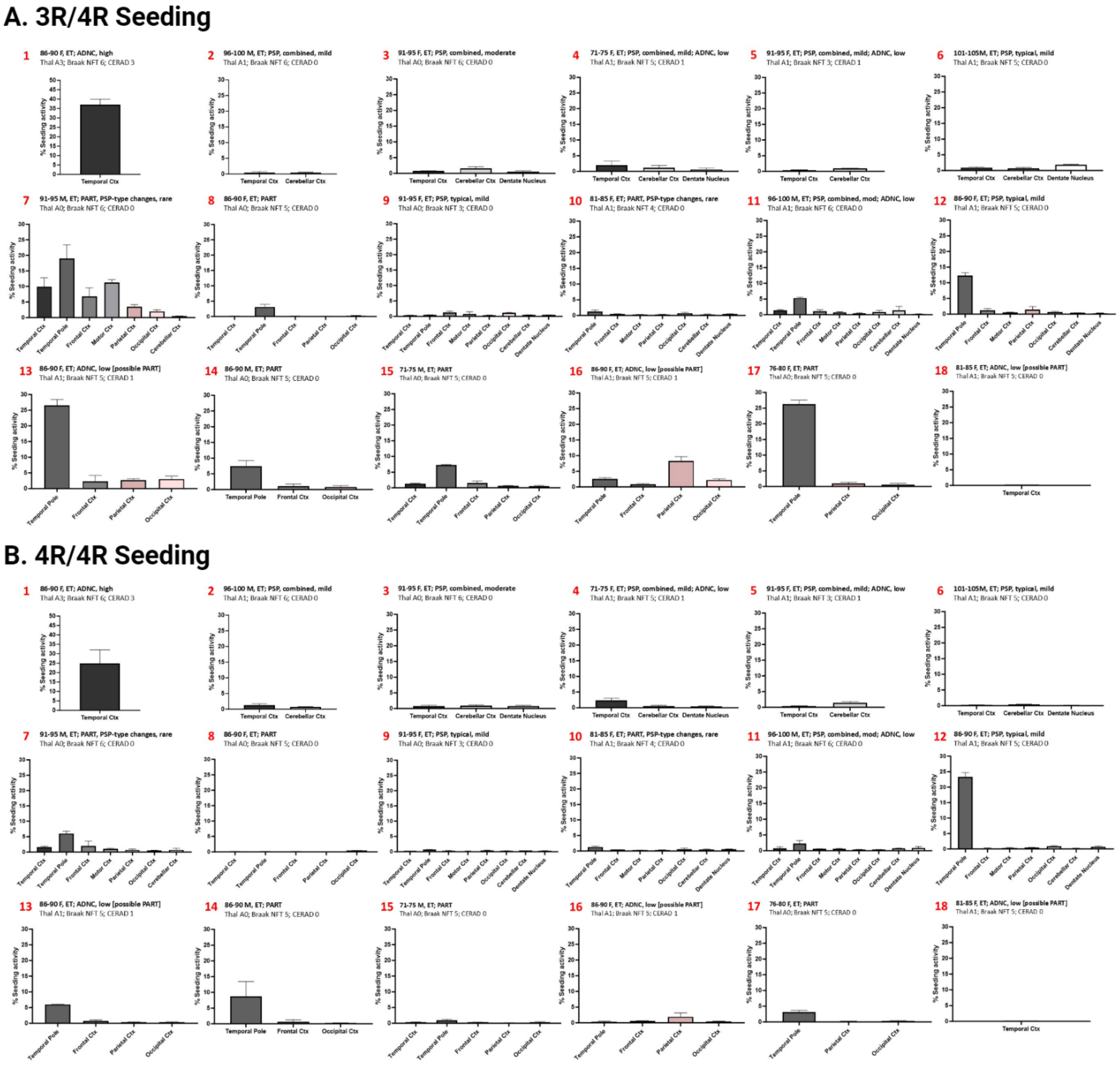


**Figure 1S: ET seeding in tau biosensor cells.** Brain homogenates from ET patients were transduced into tau RD 3R/4R (**A**) and 4R/4R (**B**) biosensor cells to measure tau seeding. Cases 1, 7, 11-15, and 17 had ≥5% seeding on the temporal cortex and temporal pole in 3R/4R biosensors. Case 16 had predominant seeding in the parietal cortex on 3R/4R biosensors. Cases 1, 7, 12-14 also had significant seeding on 4R/4R biosensors (≥5%). Cases 2, 3, 4, 5, 6, 8, 9, 10, and 18 had no/low seeding (<5%).
